# THEA: A novel approach to gene identification in phage genomes

**DOI:** 10.1101/265983

**Authors:** Katelyn McNair, Carol Zhou, Brian Souza, Robert A. Edwards

**Affiliations:** Computational Sciences Research Center, San Diego State University, 5500 Campanile Drive, San Diego, CA 92182; Lawrence Livermore National Laboratory, 7000 East Ave, Livermore, CA 94550; Department of Biology, San Diego State University, 5500 Campanile Drive, San Diego, CA 92182; Department of Computer Science, San Diego State University, 5500 Campanile Drive, San Diego, CA 92182

## Abstract

**Motivation:** Currently there are no tools specifically designed for annotating genes in phages. Several tools are available that have been adapted to run on phage genomes, but due to their underlying design they are unable to capture the full complexity of phage genomes. Phages have adapted their genomes to be extremely compact, having adjacent genes that overlap, and genes completely inside of other longer genes. This non-delineated genome structure makes it difficult for gene prediction using the currently available gene annotators. Here we present THEA (The Algorithm), a novel method for gene calling specifically designed for phage genomes. While the compact nature of genes in phages is a problem for current gene annotators, we exploit this property by treating a phage genome as a network of paths: where open reading frames are favorable, and overlaps and gaps are less favorable, but still possible. We represent this network of connections as a weighted graph, and use graph theory to find the optimal path.

**Results:** We compare THEA to other gene callers by annotating a set of 2,133 complete phage genomes from GenBank, using THEA and the three most popular gene callers. We found that the four programs agree on 82% of the total predicted genes, with THEA predicting significantly more genes than the other three. We searched for these extra genes in both GenBank’s non-redundant protein database and sequence read archive, and found that they are present at levels that suggest that these are functional protein coding genes.

**Availability and Implementation:** The source code and all files can be found at: https://github.com/deprekate/THEA

**Contact:** Katelyn McNair

## Introduction

Phages, viruses that infect bacteria, provide unique challenges for bioinformatics. There is a limit to how much DNA can be packaged in a capsid, and therefore phage genomes are generally short, typically in the range 20-100 kb. By necessity, their genomes are compact: phage genes are shorter than their bacterial homologs, are frequently co-transcribed, and adjacent open reading frames often overlap (Kang *et al.*, 2017). In a few cases, phage genes are encoded within each other (Cahill *et al.*, 2017; Summer *et al.*, 2007). In contrast, bacterial genes are longer, separated by intergenic spacers, and frequently switch strands (Kang *et al.*, 2017). There are no bioinformatics tools specifically designed to identify genes in phage genomes, so algorithms designed to identify bacterial genes are typically used (Katelyn McNair, Ramy K. Aziz, Gordon D. Pusch, Ross Overbeek, Bas E. Dutilh, and Robert Edwards, 2017). However, many of those algorithms rely on information that is not available and calculations that are not possible with short genomes. For example, there are no conserved genes in phage genomes that can be used to build training sets (Rohwer and Edwards, 2002), fewer genes means the statistics used to identify start codons are less accurate, and because many phage proteins have no homolog in the databases, similarity searches are unreliable (Roux *et al.*, 2015).

Here, we introduce a novel method for gene identification that is specifically designed for phage genomes. We presume that since phages have physical limits on their genome sizes they contain minimal non-coding DNA. This supposition allows us to take a completely novel approach to phage gene identification, tiling opening reading frames to minimize non-coding DNA bases. We treat a phage genome as a network of paths in which open reading frames are more favorable, and overlaps and gaps are less. We solved this weighted graph problem using the Bellman-Ford algorithm (Ford, 1956; Bellman, 1958).

## Materials and Methods

The first step in creating a weighted graph is finding the open reading frames. By default, we allow for three start codons (ATG, GTG, TTG), and three stop codons (TAA, TAG, TGA). The default minimum length of an ORF is 90 nucleotides. In the weighted graph, an ORF is represented as an edge between a pair of nodes: one node is the start codon and the other the stop codon. The open reading frames are then connected to nearby open reading frames, creating an edge in the graph that is either an overlap or gap (Figure 1). To save computing time, and for simplicity, we only connect ORFs within ± 300 bp of each other. When there is a very large span without an ORF, we connect ORFs on each side of the region. For each edge we calculate a weight depending on the feature type: ORF, overlap, or gap. The weight of an ORF is calculated by finding the probability of not encountering a stop codon, then taking the negative multiplicative inverse of this probability. This weight is modified by the GC frame plot score, start codon type, and ribosomal binding site score (Equation 1). Since little is currently known about the diversity of ribosomal binding sites in phages, we used the same likelihood RBS binned-system used by Prodigal (Hyatt *et al.*, 2010). We calculated a normalized global probability start codon score by counting the start codons in 2,133 phage genomes and then normalizing to one. We introduce the “GC frame plot score” (GCFP), which we calculate by identifying the frame with the maximum GC content over a 120 bp window for each codon across all ORFs that start with ATG, and then normalize to one. To account for the first possible start codon not being the correct start site, we trim off a third of the first codons, yielding a set of three position scores between 0 and 1, with 1 being the maximal frame. Then for each ORF we modify each codon’s *P*_stop_ value by the GCFP score for that particular codon. We repeat this calculation to also find the minimum GC content frame.

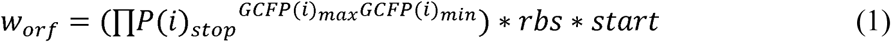

Overlap and gap weights are scaled to the ORF coding scores. Overlap weights are calculated by finding the average of the two coding weights of the ORFs in the overlap, and then exponentiating by the length of the overlap (Equation 2). If a strand switch occurs, then a strand switch penalty of 20 is added. This penalty is the multiplicative inverse of the probability of a strand switch occurring, calculated from our set of annotated genes.

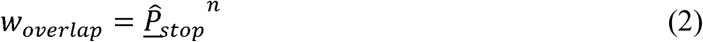

The gap weights are calculated by finding the probability of a random codon not being a stop codon across the entire genome. This probability is then exponentiated by the length of the gap, and then the positive multiplicative inverse is taken (Equation 3). If a strand switch occurs, then a penalty is added to the gap weight. For gap lengths over 300 bp we move to a linear penalty scoring calculation.

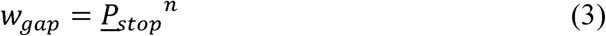

For each of the above weights, the multiplicative inverse is taken to transform the probabilities into “distances” to create a weighted graph network. The weights for open reading frames are negated; to denote these edges as favorable in the network. The Bellman-Ford algorithm is then used to find the shortest path through the network.

We compared gene identification between THEA and the three most popular gene callers: GeneMarkS, Glimmer, and Prodigal, using a set of 2,133 complete phage genomes from GenBank (Benson *et al.*, 2017). We did not include 9 Mycoplasma and Spiroplasma phages, which use an alternative genetic code. We ran THEA and each of the three alternative gene callers with default (or “phage”) parameters on each phage genome. To help mask out function, but non protein coding regions of the genomes, we used the program tRNAscan-SE to find the tRNA genes in each genome. To compare the algorithms, we counted the number of open reading frames predicted by each respective algorithm, and compared those predictions to the corresponding genes in GenBank. The source code and all files can be found at: https://github.com/deprekate/THEA

## Results

Validating phage genome annotation is frustrated by the lack of “true” phage annotations. There are few published phage proteomics studies (e.g., pmid: 26864518 25467442), and in those studies the raw proteomics data is not provided. Rather the authors only indicate which ORFs are matched, frequently using proprietary software. This precludes our ability to use proteomics data to validate gene identification in phages. Many of the current phage genome annotations in GenBank are filled with false positives. For example, in the Shiga toxin converting phages (NC_004913 and NC_004914), every ORF longer than 160 bp has been annotated as a protein coding gene. There are also abundant examples of false negatives, protein coding genes present in the genome that were not identified by the annotation software. The most obvious false negatives are genes shorter than 100bp, since this is an often-used arbitrary minimum cutoff. Small genes that do not show strong coding signals, such as shared homology to known or hypothetical genes in the databases or shared codon usage, are often excluded by other gene annotators in an effort to minimize false positives.

We calculated the number of genes predicted by each algorithm and compared them to the genes reported in GenBank. In total, we identified 225,518 genes from 2,133 phage genomes. Almost 200,000 genes were predicted by all four algorithms (Figure 1).

**Figure 1.**
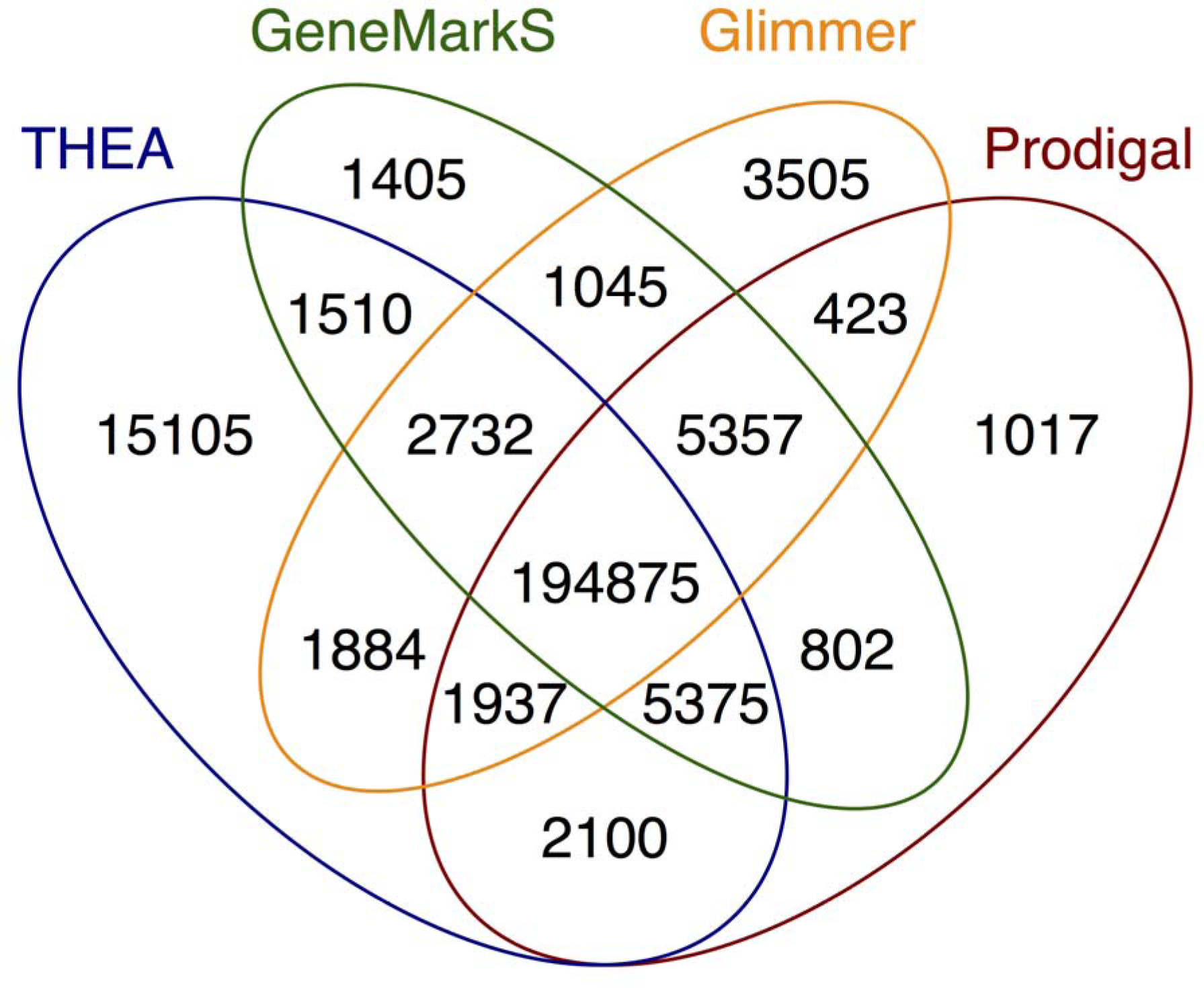
Comparison of THEA gene calls to those of three other gene annotation programs.

THEA predicted 15,105 genes that were not predicted by other gene prediction algorithms. Several lines of evidence suggest that these are phage genes are real. We first looked to see if these genes were in GenBank’s non-redundant (nr) protein database (Benson *et al.*, 2017) and found significant similarity to 23% of them (expect value < 10^-10^). This low percentage of hits is expected, since phage proteins will only have hits between 1-30% to known genes (Dutilh et al., 2012). This vast quantity of unknown phage genes is known as the “phage dark matter”. To search phage dark matter for additional hits, we performed nucleotide blast against GenBank’s Sequence Read Archive (SRA) to identify ORFs predicted by any program and presumptively non-coding ORFs not predicted by any program (i.e., all DNA stretches not interrupted by a stop codon). At the time of the search, the SRA comprised 1.3 trillion reads across 93,330 metagenomes, totalling almost 200 terabytes of data. Due to time and computational constraints, we limited the search to the first 100,000 sequences of each metagenome. We hypothesized that potential ORFs that did indeed encode a protein would occur more often in metagenomes, due to an underlying functional homology, and that potential ORFs that did not actually encode a protein would be found less frequently. For each read, we take the top hit (expect value < 10^-10^) and determine the set it is in (“ANY”, “THEA ONLY”, and “NONE”). We sum the number of hits to each set per metagenome, then divide by the number of bases in that set. We found that both the “ANY” and “THEA ONLY” gene calls occurred at similar orders of magnitude, while the “NONE” gene calls occurred much less frequently (Supplemental Fig. 1). This suggests that the gene calls that were unique to THEA, are indeed real protein encoding genes, and as such, are seen more often in phage genomes.

One of the unique features of THEA is that it is essentially reference free; aside from the GC frame plot score. Other programs, such as GeneMark and Glimmer, use hidden Markov models that require *a priori* knowledge of the composition of protein encoding genes. This is problematic when annotating phage genomes, since most potential ORFs do not have homology to any known gene. Prodigal is also unique in that it does not require a precomputed training set; however, it relies on a training set that is generated solely from the input genome. This is problematic when annotating phage genomes, since small genomes do not provide enough candidates to create a robust training set. In addition, many phage genes are horizontally transferred, and thus have different properties and signals from each other. Future versions of THEA will include the option to use these various gene properties, including hexamer frequency, codon bias, and non Shine-Dalgarno ribosomal binding site detection, and will also provide a mechanism to mask functional noncoding bases, such as those in RNAs, repeats, and *att* sites to further increase the accuracy of the gene calls.

